# Favipiravir strikes the SARS-CoV-2 at its Achilles heel, the RNA polymerase

**DOI:** 10.1101/2020.05.15.098731

**Authors:** A. Shannon, B. Selisko, NTT Le, J. Huchting, F. Touret, G. Piorkowski, V. Fattorini, F. Ferron, E. Decroly, C Meier, B. Coutard, O. Peersen, B. Canard

## Abstract

The ongoing Corona Virus Disease 2019 (COVID-19) pandemic, caused by severe acute respiratory syndrome coronavirus-2 (SARS-CoV-2), has emphasized the urgent need for antiviral therapeutics. The viral RNA-dependent-RNA-polymerase (RdRp) is a promising target with polymerase inhibitors successfully used for the treatment of several viral diseases. Here we show that Favipiravir exerts an antiviral effect as a nucleotide analogue through a combination of chain termination, slowed RNA synthesis and lethal mutagenesis. The SARS-CoV RdRp complex is at least 10-fold more active than any other viral RdRp known. It possesses both unusually high nucleotide incorporation rates and high-error rates allowing facile insertion of Favipiravir into viral RNA, provoking C-to-U and G-to-A transitions in the already low cytosine content SARS-CoV-2 genome. The coronavirus RdRp complex represents an Achilles heel for SARS-CoV, supporting nucleoside analogues as promising candidates for the treatment of COVID-19.

---

Coronaviruses (CoV) are large genome, positive-strand RNA viruses of the order *Nidovirales* that have recently attracted global attention due to the ongoing COVID-19 pandemic. Despite significant efforts to control its spread, SARS-CoV-2 has caused substantial health and economic burden, emphasizing the immediate need for antiviral treatments. As with all positive strand RNA viruses, an RdRp lies at the core of the viral replication machinery and for CoVs this is the nsp12 protein. The pivotal role of nsp12 in the viral life-cycle, lack of host homologues and high level of sequence and structural conservation makes it an optimal target for therapeutics. However, there has been remarkably little biochemical characterization of nsp12 and a lack of fundamental data to guide the design of antiviral therapeutics and study their mechanism of action (MoA). A promising class of RdRp inhibitors are nucleoside analogues (NAs), small molecule drugs that are metabolized intracellularly into their active ribonucleoside 5’-triphosphate (RTP) forms and incorporated into the nascent viral RNA by error-prone viral RdRps. This can disrupt RNA synthesis directly via chain termination, or can lead to the accumulation of deleterious mutations in the viral genome. For CoVs, the situation is complicated by the post-replicative repair capacity provided by the nsp14 exonuclease (ExoN) that is essential for maintaining the integrity of their large ~30 kb genomes^1–3^. Nsp14 has been shown to remove certain NAs after insertion by the RdRp into the nascent RNA, thus reducing their antiviral effects^4–6^. Despite this, several NAs currently being used for the treatment of other viral infections have been identified as potential anti-CoV candidates^7–9^. Among these is the purine base analogue T-705 (Favipiravir, Avigan, Extended data Fig. 1a,b) that has broad-spectrum activity against a number of RNA viruses and is currently licensed in Japan for use in the treatment of influenza virus^10^. Clinical trials are currently ongoing in China, Italy, and the UK for the treatment of COVID-19, although a precise MoA of the drug against CoVs is lacking.

We infected Vero cells with CoV-SARS-2 in the presence or absence of 500 μM T-705 and performed deep sequencing of viral RNA. A 3-fold (P<0.001) increase in total mutation frequencies is observed in viral populations grown in the presence of the drug as compared to the no-drug samples (Fig. 1). Similar to previous findings with influenza^11^, Coxsackie B3^5^ and ebola^12^ viruses, a 12-fold increase in G-to-A and C-to-U transition mutations is observed, consistent with T-705 acting predominantly as a guanosine analogue. The increase in the diversity of the virus variant population suggests that once incorporated into viral RNA, T-705 is acting as a mutagen capable of escaping the CoV repair machinery. Interestingly, the SARS-CoV-2 genome has an already low cytosine content of ~17.6% and T-705 treatment may therefore place additional pressure on the already unbalanced CoV nucleotide content. Associated with this increase in mutation frequency, T-705 has an antiviral effect on SARS-CoV-2, as illustrated by a reduction in virus-induced cytopathic effect, viral RNA copy number, and infectious particle yield. Altogether these observations show that the mutagenic effect induced by T-705 is, at least in part, responsible for the inhibition of the replication.

**Fig. 1:**
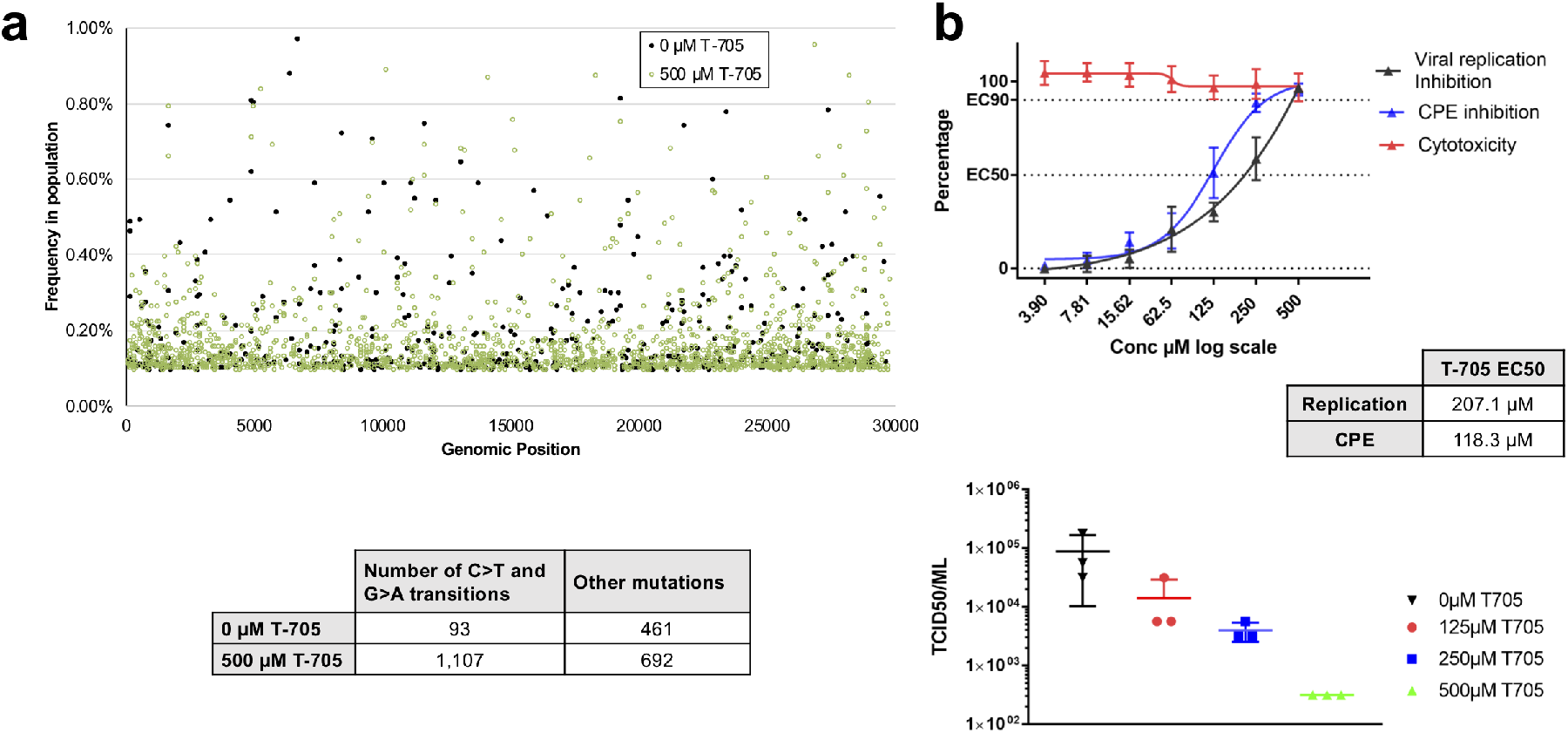
Antiviral effects of T-705 on SARS-CoV-2. **a,** *In vitro* effects of T-705 on SARS-CoV-2. Distribution of the mutations along the SARS-CoV-2 genome and number of mutations observed in presence or absence of T-705. A 3-fold increase in the presence of the drug is observed (P<0.001, Pearson’s Chi-squared test with Yates’ continuity correction). **b,** Quantification of the antiviral effect of T-705 by genome copy number, virus-mediated CPE, and number of infectious particles.

To determine the efficacy and MoA of T-705 against SARS-CoV we first characterized nsp12 primer-dependent activity using traditional annealed primer-template (PT) and self-priming hairpin (HP) RNAs that may confer additional stability on the elongation complex (Extended data Fig. 1c). Consistent with prior findings, nsp12 alone is essentially inactive^13,14^ and RNA synthesis requires the presence nsp7 and 8 cofactors whose stimulatory effect is enhanced by linking them as a nsp7L8 fusion protein^6,15^. Structures of nsp12 show a four-component complex with a nsp7/nsp8 heterodimer and an additional nsp8 monomer^16,17^ and accordingly we found that addition of supplementary nsp8 to the nsp12:nsp7L8 complex further increases activity (Extended data Fig. 2). The resulting nsp12:7L8:8 complex is highly active on both PT and HP RNAs, with reactions containing 0.2 μM of each substrate and 1 μM nsp12 showing comparable initiation rates, with a rapid burst phase resulting in >50% primer consumption followed by remaining primer use over a period of a few minutes (Fig. 2a, Extended data Fig. 3). Interestingly however, the apparent processivity for the two substrates differs substantially. When provided with an annealed PT pair, intermediate products account for ~60% of the total lane intensity across all timepoints suggesting distributive polymerase activity (Fig. 2a, Extended data Fig. 4). In contrast, extension of a HP substrate has few intermediate products, indicating a processive elongation mode. This pattern is consistently observed across RNA substrates of different lengths, showing that the distributive PT mode does not convert into a processive state within a 30-nucleotide long template. Notably, the *Nidovirales* RNA replication/transcription scheme involves precise recombination-like events to generate subgenomic RNAs through a discontinuous mechanism along the ~30,000 nt genome^18^. The different processivities and complex stabilities we observe may be connected to these peculiar RNA synthesis events.

**Fig. 2:**
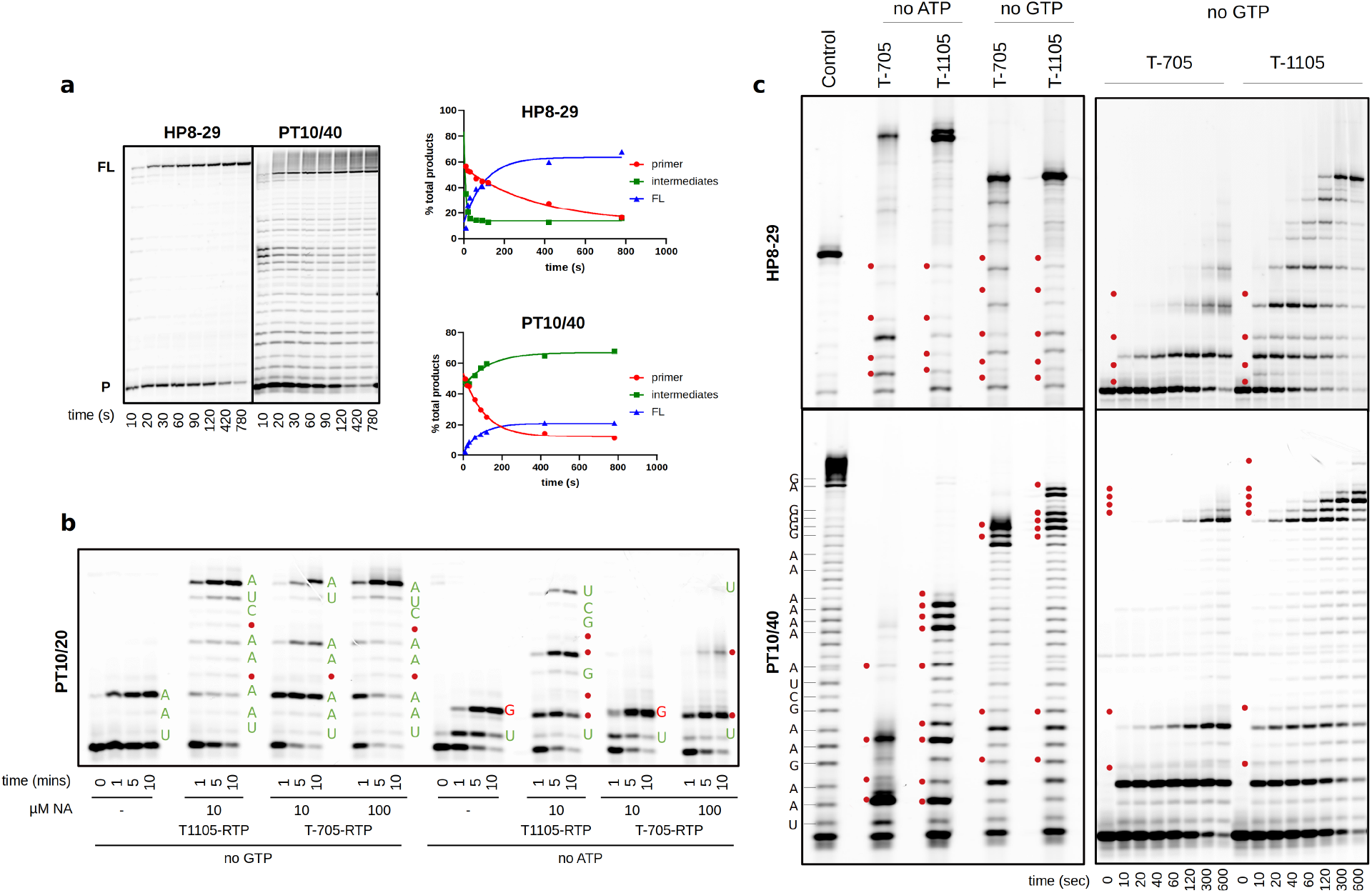
Polymerization modes of SARS-CoV nsp12 and incorporation of nucleotide analogues. **a,** Steady-state data showing extension of hairpin and primer-template RNA substrates by the nsp12:7L8:8 complex. Graphs show the fraction of primer, intermediate and full-length products over time. Additional extension products beyond the full-length observed on the PT substrate are attributed to nsp8 RNA 3’-terminal adenylyltransferase activity^40^. **b,c,** Incorporation of nucleoside analogues T-705 and T1105 with different RNA substrates in the absence of GTP or ATP. Red dots show analogue incorporation sites at the omitted nucleotide positions. Incorporation of correct nucleotides shown in green and mismatches in red. **c,** Left panel shows products obtained with 100 μM each analogue after 5 minutes. Right panel shows time-course in the presence of 10 μM of each analogue and absence of GTP. Only partial site assignments were possible with the hairpin substrate due to drastic differences in RNA migration attributed to changes in residual RNA structure following analogue incorporation.

We sought to determine the extent to which the RdRp complex could incorporate the NA inhibitors T-705 and T-1105 (a non-fluorinated T-705-related analogue, Extended data Fig. 1a,b) into RNA. T-1105 shows improved potency against influenza virus in certain cell lines and is more stable than T-705 in the RTP form used in enzyme assays^19–21^. The MoA for these compounds is currently controversial. T-705 has been shown to act through lethal mutagenesis for several viruses, predominantly by competing with guanosine to cause transition mutations^11,22–25^. However, two separate studies support an antiviral effect mediated by chain termination, with incorporation of either a single or two consecutive T-705 molecules blocking further extension by influenza polymerase^26,27^.

For the nsp12 complex, omission of ATP and/or GTP from elongation reactions results in rapid incorporation of T-705 and T-1105 at multiple sites with both substrates (Fig. 2b,c). In reactions with 10 μM T-1105-RTP, multiple incorporation events are seen within 10 seconds, while T-705-RTP is less efficient. Neither compound is incorporated in the place of cytosine or uracil, clearly showing they function as purine analogues (Extended data Fig. 5). For the processive HP complex, efficient incorporation and elongation occurs opposite both uracil and cytosine, resulting in rapid accumulation of full-length products, advocating for lethal mutagenesis as the MoA for these compounds (Fig. 2c). Interestingly, the PT substrate reveals a striking difference in the MoA depending on whether the analogues are incorporated in place of guanine or adenine. Opposite uracil, both are rapidly incorporated but further extension is slow and inefficient, leading to an accumulation of RNA products consistent with a non-obligate chain termination mechanism. The presence of a non-natural 3’ nucleotide analogue may destabilize the complex and promote template dissociation. In contrast, opposite cytosine there is a stall before each analogue incorporation step, but elongation past the analogue is rapid, even at consecutive sites. More full-length product is observed in these no-GTP experiments, suggesting both compounds are more efficient as guanosine analogues. A similar observation has been made for the poliovirus RdRp that efficiently incorporated and bypassed T-1105 opposite cytosine but was prone to pausing opposite uridine. These pause events were attributed to RdRp backtracking, which was therefore assigned the primary cause of the inhibitory activity^28^. We conclude that for nsp12 the T-705/T-1105 MoA is a combination of chain termination, slowed RNA synthesis and lethal mutagenesis, dictated by the structural and functional properties of both the polymerase and the RNA.

The reactions performed without ATP also showed detectable amounts of GTP:U mismatch products, allowing us to make a direct comparison of the T-1105 incorporation rate with that of a natural GTP:U mismatch (Extended data Fig. 6). Reactions using 50 μM GTP and 1 μM T-1105-RTP show ~5-fold more T-1105:U produced relative to GTP:U. Considering the concentration difference, T-1105 incorporation is calculated to be ~250-fold more efficient than the native GTP mismatch, the most common naturally occurring transition mutation. These high incorporation rates likely overwhelm the CoV error-correcting mechanisms, consistent with our infectious virus data, although it remains to be determined if nsp14 can excise T-705/T-1105, as is the case for ribavirin and 5-fluorouracil^4,6,29^.

We carried out pre-steady state rapid-quench experiments to further understand the molecular basis of nsp12 elongation rates and fidelity. We initially attempted to form stalled elongation complexes, as is commonly done for the structurally related picornaviral RdRps ^30–32^ but nsp12 complexes showed half-lives of only ~1 minute across a range of conditions and cofactor stoichiometries (Extended data Fig. 7). The significant rapid burst phase, however, allowed characterization of elongation rates under single-turnover conditions using millisecond timescale EDTA quench-flow (Fig. 3 and Extended data Fig. 8). Experiments were performed at three NTP concentrations based on estimated steady-state K_d_ values of 3-15 μM. CTP was omitted to allow elongation by only 7 and 8 nucleotides on PT and HP substrates respectively, while avoiding end-effects that can slow elongation rates. At 17 μM NTP we observe multiple incorporation steps within a mere 10-20 ms and formation of the +7/8 products by 100 ms (Fig. 3). Analysis of the data using a minimal model of identical nucleotide incorporation steps yields elongation rates of 66±1.3 and 95±8 nt/s for the PT and HP substrates at 50 μM NTP at 22 °C (Extended data Table 1). Fitting the rate concentration dependence yields maximal elongation rates of 90±4 nt/s on annealed primer-templates and 150±30 nt/s on hairpin templates (Fig. 3d).

**Fig. 3.**
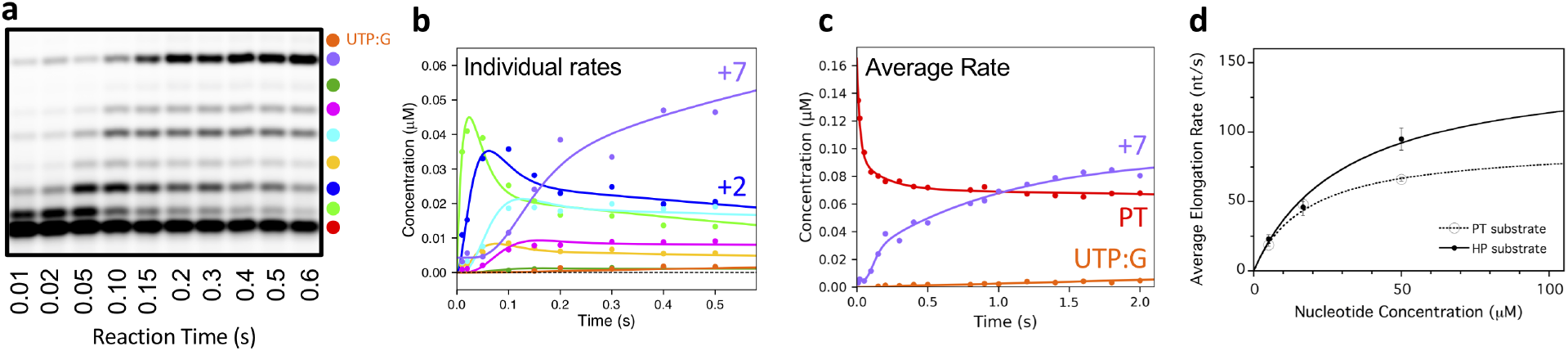
Pre steady-state elongation by nsp12-7L8-8 complex. **a,** EDTA quench-flow data showing rapid elongation to +7 products on the PT10/20 substrate in the absence of CTP. **b,c,** Fitting of the quench-flow data to models with discrete rates for each elongation step (**b,** color key in **a**) or with a single average incorporation rate for each step (**c**). **d,** Nucleotide concentration dependence yielding maximal average elongation rates of 150 ± 30 nt/s on the HP substrate and 91 ± 4 nt/s on the PT substrate.

These data reveal that the SARS-CoV nsp12 is the fastest viral RdRp known, with rates significantly faster than the 5-20 nt/s observed for picornaviral polymerases at room temperature^32–34^ and 4-18 nt/s for hepatitis C and dengue virus polymerases at 30 and 37° C^35,36^. Based on its structure, nsp12 is expected to use the palm domain based active site closure mechanism that is unique to viral RdRps^31^ and which exhibits a 3-to-5 fold rate increase between 22 °C and 37 °C^33,34^, suggesting nsp12 can elongate at a staggering 600-700 nt/s at physiological temperatures.

Such a fast viral RdRp is consistent with the need to rapidly replicate ~30,000 nt long RNA genomes, but raises questions as to how fidelity of nucleotide incorporation is impacted. In our data, we consistently observe nucleotide misincorporations and subsequent elongation on templates where nsp12 should stall due to the lack of CTP (Extended data Fig. 2b, 3, 9). In contrast, RdRps that form highly stable elongation complexes show limited or no read-through products under comparable conditions^30,35,37^. The efficiency of the nsp12 bypass is UTP concentration dependent (Extended data Fig. 3), indicative of uracil misincorporation opposite a templating guanosine (UTP:G). The UTP:G elongation product is also observed in the PT quench-flow data at reaction times as short as 100 ms, yielding a misincorporation rate of ~0.2 nt/s (Extended data Table 1). This is only 400–500-fold lower than the cognate UTP:A rate measured in the same experiment, more than one order of magnitude less accurate than the generally admitted 10^-4^-10^-6^ error rate of viral RdRps^38^.

The molecular basis for fast and low fidelity replication by nsp12 is not yet known, but a comparison of RdRp structures reveals that a key NTP interaction is absent in CoV enzymes (Fig. 4). Viral RdRps use an electrostatic interaction with an arginine residue in motif F to position the NTP during catalysis^31^. For most positive strand RNA virus polymerases, this arginine is stabilized by a salt bridge to a glutamic acid residue, also from motif F. Notably, the CoV nsp12 has an alanine in place of this glutamate (A547) and as a result, the arginine (R555) is not rigidly anchored above the active site (Fig. 4a)^31,39^. This flexibility could allow catalysis to occur with a relaxed triphosphate positioning, decreasing fidelity by lowering the requirement for strict Watson-Crick base pairing in the active site. Interestingly, the CoV and other large-genome nidoviral polymerases also have an SDD rather than a more typical GDD sequence in the palm domain of motif C (Fig. 4b). This serine residue can hydrogen bond with the 2’ hydroxyl of the priming nucleotide, theoretically stabilizing the position of the primer in the active site and compensating for the less rigid interactions with the bound NTP. These unique features have likely played a central role in genome expansion and stability by providing a fast RNA synthesis machinery whose inaccuracy is mitigated by the presence of an RNA repair exonuclease. Our data demonstrate that nucleoside analogues are pertinent candidates for the treatment of COVID-19. Favipiravir, with its already defined safety profile and mode of action, may well find a place as an anti-RdRp component in combination therapies targeting coronaviruses.

**Fig. 4.**
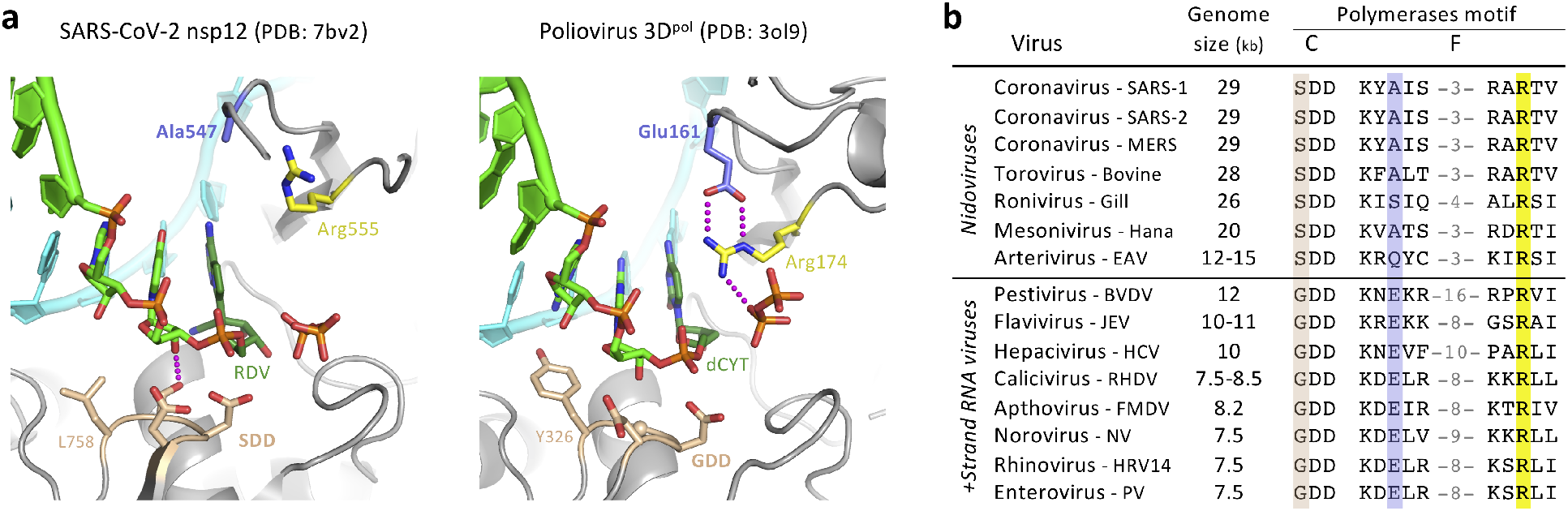
Comparison of viral polymerase active site structures and sequences. **a,** Analogous post-catalysis, pre-translocation structures of CoV nsp12 and poliovirus 3Dpol after incorporation of remdesivir (RDV) and deoxycytosine (dCYT), respectively. The CoV polymerases have replaced a motif F glutamate residue (Glu161) with an alanine (Ala547), removing a highly conserved interaction that positions the motif F arginine for interactions with the NTP and pyrophosphate. **b,** Alignment of representative viral RdRp sequences showing the large genome nidoviruses have a SDD instead of GDD sequence in the palm domain motif C (tan) and alanine, serine, or glutamine in place of the aforementioned glutamate (blue) found in other positive strand RNA viruses. The NTP-interacting arginine (yellow) is conserved, but the overall length of the motif F loop is shorter in the nidoviruses (numbers reflect omitted residues).

## Supporting information

ExtendedData_and_figures

## Acknowledgments

We thank Magali Gilles for excellent technical support as well as Prs. C. Drosten and F. Drexler for providing the SARS-CoV-2 through EVA-GLOBAL (European Union’s Horizon 2020 programme, GA 871029). This work was supported by the Fondation pour la Recherche Médicale (Aide aux équipes), the SCORE project H2020 SC1-PHE-Coronavirus-2020 (grant#101003627) to BCa, Inserm through the REACTing initiative (REsearch and ACTion targeting emerging infectious diseases) to BCa and BCo, National Institutes of Health grant AI059130 to OP, and a grant from DZIF (German Center for Infection Research) to JH and CM.

## Author contributions

BS/NTTL/VF performed experiments and analyzed data, JH/CM performed experiments, FT/GP/BCo performed experiments and analyzed data, FF/ED analyzed data, AS/OP/BCa designed experiment, performed experiments, analyzed data and wrote manuscript.

## Materials & Correspondence

Correspondence to: either

## Materials and Methods

### Nucleotide substrates

T-1105-ribose-TP (RTP), the 5’-triphosphate of the non-fluorinated derivative of T-705-ribose, was synthesized as described previously^21^ following the iterative route from mono-via di-to triphosphate with minor modifications. To overcome low solubility of T-1105-ribonucleotides in organic solvent, the 2’- and 3’-acetyl groups were kept until the final TP-synthesis and were then cleaved by treatment with base. This reaction sequence resulted in an increased yield of up to 90% (for di-) and 46% (for triphosphate), nearly double of previously reported. T-705-RTP was obtained from Toronto Research Chemicals. Lyophilized aliquots were resuspended in TE buffer pH 7.0 and used the same day. Stability can be verified by measuring Abs at 372 nm and 350 nm for T-705-RTP and T-1105-RTP respectively. Other NTPs were purchased from GE Healthcare.

### Synthetic Oligonucleotides

Primer-template (PT) pairs were purchased from Biomers (HPLC grade). RNA oligonucleotides ST20 (20-mer) and LS1 (40-mer), corresponding to the 3’-end of the SARS-CoV genome (excluding the poly A tail) were used as templates, annealed to a 5’ cy5 labeled SP10 primer (10-mer) corresponding to the 5’-end of the anti-genome. For simplicity, these were named PT10/20 and PT10/40 throughout the manuscript. Annealing was performed by denaturing primer:template pairs (molar ratio of 1:1.5) in 110 mM KCl at 70° C for 10 min, then cooling slowly to room temperature over several hours. Hairpin RNAs were synthesized by Integrated DNA Technologies (Coralville, IA), resuspended at 100 uM concentration in 50 mM NaCl, 5 mM MgCl_2_, 10 mM Tris (pH 8.0) and heated to 95°C for 15 minutes before snap cooling on ice.

### Expression and Purification of SARS-CoV proteins

All SARS-CoV proteins used in this study were expressed in *Escherichia coli* (*E. coli*), under the control of T5 promoters. Cofactors nsp7L8 and nsp8 alone were expressed from pQE30 vectors with C-terminal and N-terminal hexa-histidine tags respectively. TEV cleavage site sequences were included for His-tag removal following expression. The nsp7L8 fusion protein was generated by inserting a GSGSGS linker between nsp7- and nsp8-coding sequences. Cofactors were expressed in NEB Express C2523 (New England Biolabs) cells carrying the pRare2LacI (Novagen) plasmid in the presence of Ampicillin (100 μM/mL) and Chloramphenicol (17 μg/mL). Protein expression was induced with 100 μM IPTG once the OD_600_ = 0.5–0.6, and expressed overnight at 17°C. Protein was purified first through affinity chromatography with HisPur Cobalt resin (Thermo Scientific), with a lysis buffer containing 50 mM Tris-HCl pH 8, 300 mM NaCl, 10 mM Imidazole, supplemented with 20 mM MgSO4, 0.25 mg/mL Lysozyme, 10 μg/mL DNase, 1 mM PMSF, with lysis buffer supplemented with 250 mM imidazole. Eluted protein was concentrated and dialyzed overnight in the presence of histidine labeled TEV protease (1:10 w/w ratio to TEV:protein) for removal of the protein tag. Cleaved protein was purified through a second cobalt column and protein was purified through size exclusion chromatography (GE, Superdex S200) in gel filtration buffer (25 mM HEPES pH 8, 150 mM NaCl, 5mM MgCl_2_, 5 mM TCEP). Concentrated aliquots of protein were flash-frozen in liquid nitrogen and stored at −80 °C. A synthetic, codon-optimized SARS-CoV nsp12 gene (DNA 2.0) bearing C-terminal 8His-tag preceded by a TEV protease cleavage site was expressed from a pJ404 vector (DNA 2.0) in *E. coli* strain BL21/pG-Tf2 (Takara). Cells were grown at 37°C in the presence of Ampicillin and Chloramphenicol until OD600 reached 2. Cultures were induced with 250 μM IPTG and protein expressed at 17°C overnight. Purification was performed as above in lysis buffer supplemented with 1% CHAPS. Two additional wash steps were performed prior to elution, with buffer supplemented with 20 mM imidazole and 50 mM arginine for the first and second washes respectively. Polymerase was eluted using lysis buffer with 500 mM imidazole and concentrated protein was purified through gel filtration chromatography (GE, Superdex S200) in the same buffer as for nsp7L8. Collected fractions were concentrated and supplemented with 50% glycerol final concentration and stored at −20 °C.

### Steady-state elongation complex reactions

Nsp12, nsp7L8 and nsp8 were mixed just prior to each experiment at a 1:3:3 molar ratio (unless otherwise stated) and preincubated on ice for 10 mins. The protein complex was subsequently mixed with the RNA pre-mix (20 mM HEPES pH 7.5, 50 mM NaCl, 5 mM MgCl_2_) containing either a single RNA substrate, or both HP and PT RNAs at equimolar ratios. Reactions were initiated with 50 μM (final concentration) of all four NTPs, or without CTP for partial elongation reactions. Final reaction concentrations were 1 μM nsp12, 0.2 μM RNA. Reactions were quenched at indicated timepoints with 5X volume of FBD stop solution (formamide, 10 mM EDTA). To verify that activity was not due to either nsp7L8 or nsp8, which have been shown to contain primase-like noncanonical RdRp activities^40–43^ assays were run in the absence of nsp12. The NTP Kd was estimated using the same setup described here, but with final (equimolar) concentration of ATP, GTP and UTP ranging from 0.78 – 100 μM (2-fold dilution series). The time course of product formation was fit to single exponential equation for each concentration of NTP to give the observed rate constant (kobs). Observed rates were subsequently plotted against NTP concentration, and the data was fit via hyperbolic regression to give the equilibrium dissociation constant (Kd) and the maximum rate constant for incorporation of NTPs (kpol).

### Formation of stalled elongation complex

Attempts to form a stalled elongation complex were performed with HP8-22 RNA with varied ratios of nsp12:nsp7L8:nsp8 (using a constant nsp12 concentration 0.8 μM). Protein, HP8-22 RNA (0.2 μM) and 50 μM final concentrations of GTP and ATP were incubated for 5 min at room temperature to allow formation of a +4-lock stalled complex. Reactions were diluted 1:2 in high salt to prevent RNA rebinding, and chased at various timepoints (1 – 50 minutes) with 50 μM all NTPs. Chase reactions were quenched after 30 sec in FBD. Stability half-life was calculated from the ratio of full-length (FL) product produced relative to the total amount of +4-lock + FL product at each timepoint. Half-life was obtained by fitting the data through a single exponential.

### Pre-steady state quenched-flow kinetics

Experiments were carried out in a Bio-Logic QFM-300 rapid chemical quench-flow apparatus that controls reaction time by flow rate through a 2.5 uL chamber between the reaction mixer and the quencher mixer. The nsp12-nsp7L8-nsp8 complex was assembled by first preincubating the proteins on ice for 10 minutes at a 1:3:3 molar ratio, then adding this to SP10/20 and HP8-29 hairpin RNA in reaction buffer (20 mM HEPES pH 7.5, 5mM MgCl_2_, 50 mM NaCl) and further incubated at room temperature for 15 minutes prior to beginning the rapid-quench experiments, which took ~10 minutes to complete a set of 22 different reaction times. Reactions were initiated by mixing equal volumes (18 ul each) of the protein-RNA complex with NTP solutions containing equimolar concentrations of ATP, GTP and UTP, and then quenched with 18 uL of 200 mM EDTA. Final concentrations in the reactions were 1 uM nsp12, 0.2 uM primer-template RNA, 0.2 uM hairpin RNA, and 50, 16.7 or 5.6 uM NTP, and the final EDTA concentration after quenching was 67 mM.

### Product Analysis

Quenched reactions were mixed 1:1 with FBD loading dye and heated for 10 min at 70°C, and cooled on ice for 2 min before analysis on 17 – 20% polyacrylamide, 7 M Urea TBE gels. Gels were run at a constant 65 W using Sequi-Gen GT Systems from Bio-Rad or vertical electrophoresis systems from CBS Scientific and visualized using an Amersham™ Typhoon™ Biomolecular Imager (GE Healthcare). The intensity of each band was quantified using the ImageQuant software (GE Healthcare)/Image Gauge (Fuji) and/or using ImageJ as implemented in the Fiji package, with background subtraction. Product yield was determined by dividing the intensity of the product by total intensity of the product + remaining primer, and multiplying by the input concentration of RNA. For the rapid quench data, the program PeakFit (Systat Software) was used to fit gel lane profiles to a set of gaussian peaks and the fractional area contained within each peak was multiplied by the RNA concentration in the experiment to calculate the amount of each elongation species as a function of reaction time.

### Pre-steady state kinetic data analysis

Rapid quench date were analysed using the program KinTek Explorer^44^ to model the reactions as a series of seven (PT substrate) or eight (HP substrate) irreversible nucleotide addition steps. Data from each NTP concentration series were fit independently (not globally) to obtain observed rates using either a model with i) a single average rate or ii) individual rates for each nucleotide addition step (fig S8). Modeling the remaining intermediate species due to low processivity elongation using formal RNA dissociation and rebinding steps (at rates comparable to non-burst phase primer utilization) was a not successful, suggesting the nsp12-RNA complex exists in some form of inactive state. Low processivity elongation was instead accommodated in the kinetic model by adding a reversible inactivation/reactivation equilibrium step for each elongation product, allowing intermediate products to depart from the immediate processive pathway toward full length product formation, but then rejoin the pathway for further elongation. The rate constants for these steps were shared among all intermediates species as there was insufficient data to fit them individually.

The single average rate models were fit using data from primer loss and the expected fulllength (+7 or +8) product, but not the intermediate species. Note that for the PT template we observed detectable amounts of additional +8 and +9 bands at the higher 16.7 and 50 μM NTP concentrations and therefore included them in the model. This is presumably due to a UTP:G mismatch to yield the +8 product in the absence of CTP, followed by a cognate ATP:U addition that is slow because it is priming on a mismatched base pair. For the HP substrate, we were unable to definitively identify the gel migration bands for the +6 and +7 species and therefore did not model individual rates for these incorporation events. Data from the HP RNAs are can be challenging to analyze because the high thermodynamic stability of the folded RNA structure can make it difficult to fully denature the helix during electrophoresis. This leads to a mixture of species that transition from denatured single stranded RNA for the initial species to more compact and faster migrating duplexes for longer elongation products. We were therefore unable to reliably analyze quench flow data from longer hairpin products.

### NTP-analog incorporation

Enzyme mix (20 μM nsp12, 60 nsp7L8, 60 μM nsp8) in complex buffer (25 mM HEPES pH 7.5, 150 NaCl, 5 mM TCEP, 5 mM MgCl_2_) was incubated 10 min on ice and then diluted with reaction buffer (20 mM HEPES pH 7.5, 50 mM NaCl, 5 mM MgCl_2_) to 4 μM nsp12 (12 nsp7L8 and 8). The resulting enzyme complex was mixed with the same volume of 0.8 μM primer/ template (P/T) with or without 0.8 μM hairpin (HP) carrying 6-FAM or Cy5 fluorescent labels at their 5’ ends in reaction buffer, and incubated at 25°C (or 30°C as given in Figure legends) for 10 min. Reactions were then started with the same volume of 100 μM NTPs (leaving out one or two as given in Figure legends) and TE buffer or NTP-analogs (T-1105-RTP or T-705-RTP) in reaction buffer. Final concentrations in the reactions were 1 μM nsp12 (3 μM nsp7L8 and 8), 0.2 μM P/T (and 0.2 μM HP), 50 μM NTPs and the given concentrations of NTP-analogs. Samples of 8 μl were taken at given time points and mixed with 40 μl of formamide containing 10 mM EDTA. Ten-μl samples were analyzed by denaturing PAGE (20 % acrylamide, 7 M urea, TBE buffer); and product profiles visualized by a fluorescence imager (Amersham Typhoon).

### In vitro infection assays

Cell line: VeroE6 (ATCC CRL-1586) cells were grown in minimal essential medium (Life Technologies) with 7.5% heat-inactivated fetal calf serum (FCS), at 37 °C with 5 % CO_2_ with 1 % penicillin/streptomycin (PS, 5000 U.mL^-1^ and 5000 μg.mL^-1^respectively; Life Technologies) and supplemented with 1% non-essential amino acids (Life Technologies).

Virus strain: SARS-CoV-2 strain BavPat1 was obtained from Pr. Drosten through EVA GLOBAL (https://www.european-virus-archive.com/). Virus stock was prepared as previously described^45^. All experiments were conducted in a BSL3 laboratory.

Antiviral experiments: For EC50 and CC50 determination, one day prior to infection, 5×10^4^ VeroE6 cells were seeded in 100 μL assay medium (containing 2.5 % FCS) in 96 well plates. The next day, seven 2-fold serial dilutions of T-705 (500 μM to 3.9 μM in triplicate) were added to the cells (25 μL/well, in assay medium). Four virus control wells were supplemented with 25 μL of assay medium. After 15 min, 25 μL of a virus mix diluted in medium was added to the wells. The amount of virus working stock used was calibrated prior to the assay, based on a replication kinetics, so that the replication growth is still in the exponential growth phase for the readout as previously described^46,47^. Four cell control wells (*i.e.* with no virus) were supplemented with 50 μL of assay medium. Plates were incubated for 2 days at 37 °C prior to quantification of the viral genome by real-time RT-PCR. RNA extraction and viral RNA quantification was performed has previously described^45^. The 50% and 90% effective concentrations (EC50, EC90; compound concentration required to inhibit viral RNA replication by 50% and 90%) were determined using logarithmic interpolation as previously described^46^. For the evaluation of CC50 (the concentration that reduces the total cell number by 50%), the same culture conditions were set as for the determination of the EC50, without addition of the virus, then cell viability was measured using CellTiter Blue^®^ (Promega). Cell supernatant media were discarded and CellTiter-Blue^®^ reagent (Promega) was added following the manufacturer’s instructions. Plates were incubated for 2 hours prior recording fluorescence (560/590nm) with a Tecan Infinite 200Pro machine. From the measured OD_590_, CC50 was determined using logarithmic interpolation. For EC50 determination using CPE inhibition, cells and T-705 were prepared as previously described (EC50 determination). Eight virus control wells were supplemented with 25 μL of assay medium and eight cell control were supplemented with 50 μL. After 15 min, 25 μL of a virus mix diluted in 2.5 % FCS-containing medium was added to the wells at MOI 0.002. Three days after infection, CPE were assessed using CellTiter-Blue^®^ reagent (Promega). For the infectivity test cells and compound were prepared as described above (EC50 determination) with only three 2-fold serial dilutions of T-705 (125μM to 500μM, in triplicate). Three virus control wells were supplemented with 25 μL of assay medium. The experiment was conducted as previously described for the EC50 determination. At day 3 the supernatant was collected and each triplicate was titrated by measuring the 50 % tissue culture infectivity dose (TCID_50_); briefly, 3 replicates were infected with 100 μL of tenfold serial dilutions of the previous supernatant, and incubated for 3 days. CPE was measured by CellTiter-Blue^®^ reagent (Promega) and TCID50 was calculated and expressed as TCID_50_/mL based on the Reed and Muench *(43).* Antiviral experiments data were analyzed using GraphPad Prism 7 software (Graph pad software). Graphical representations were also performed on GraphPad Prism 7 software.

### Sequence analysis

Eight overlapping amplicons were produced from the extracted viral RNA using the SuperScript *IV* One-Step *RT-PCR System* (Thermo Fisher Scientific) and specific primers (supplemental data). PCR products were pooled at equimolar proportions. After Qubit quantification using Qubit^®^ dsDNA HS Assay Kit and Qubit 2.0 fluorometer (ThermoFisher Scientific) amplicons were fragmented by sonication in ~200bp long fragments. Libraries were built adding barcodes for sample identification to the fragmented DNA using AB Library Builder System (ThermoFisher Scientific). To pool the barcoded samples at equimolar ratio a quantification step by real time PCR using Ion Library TaqMan™ Quantitation Kit (Thermo Fisher Scientific) was realised. An emulsion PCR of the pools and loading on 530 chip was realised using the automated Ion Chef instrument (ThermoFisher). Sequencing was performed using the S5 Ion torrent technology v5.12 (Thermo Fisher Scientific) following manufacturer’s instructions. Consensus sequence was obtained after trimming of reads (reads with quality score <0.99, and length <100pb were removed and the 30 first and 30 last nucleotides were removed from the reads) mapping of the reads on a reference (determined following Blast of De Novo contigs) using CLC genomics workbench software v.20 (Qiagen). A *de novo* contig was also produced to ensure that the consensus sequence was not affected by the reference sequence. Quasi species with frequency over 0.1% were studied.

