## ExtendedData_and_figures for "Favipiravir strikes the SARS-CoV-2 at its Achilles heel, the RNA polymerase"

**This PDF file includes:**

Extended data Figs. 1 to 9
Extended data Table 1

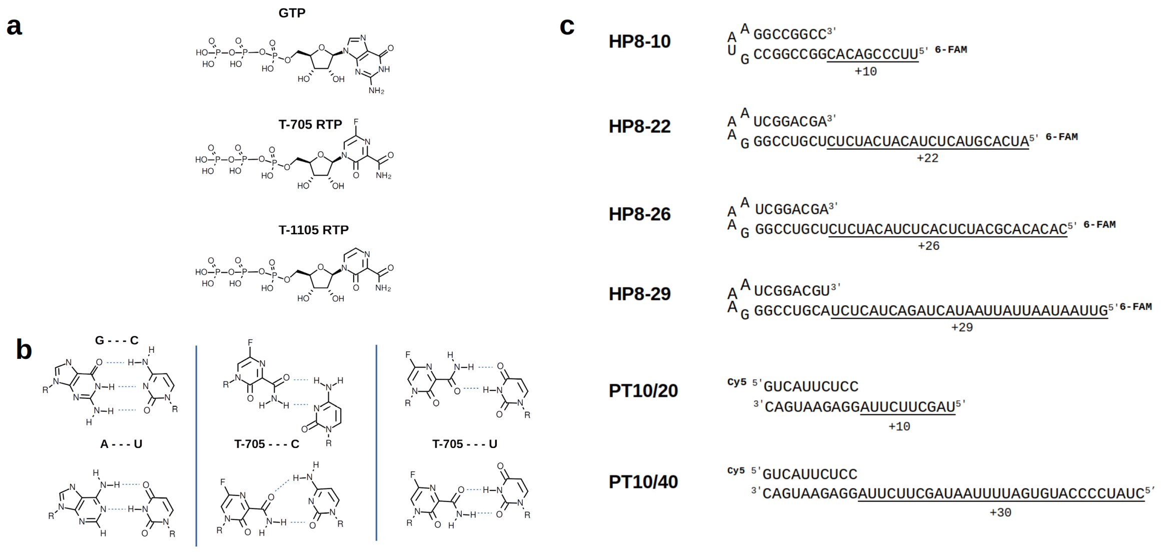

**Extended data Fig. 1. Nucleoside analogues and RNA substrates. a,** Chemical structures of the nucleotide guanosine 5'-triphosphate (GTP), the chain terminator T-705 ribofuranosyl 5'-triphosphate (T-705-RTP), and its non-fluorinated analogue (T-1105-RTP). **b,** Schematics of standard base pairing guanine – cytidine and adenine – uridine (left) versus ambiguous base pairing of Favipiravir (T-705) opposite to a cytidine (central) or a Uridine (right) RNA template. **c,** Hairpin RNAs (HP) are labelled at the 5' end with 6-Carboxyfluorescein (6-FAM). Nomenclature of HP8-X refers to the length of the duplex stem (8) and length of the overhanging, single-stranded templating strand for extension (X). The primer of the primer-template (PT) substrates are 5' labelled with cyanine-5 (Cy5). Numbers refer to the length of the primer and template strand respectively.

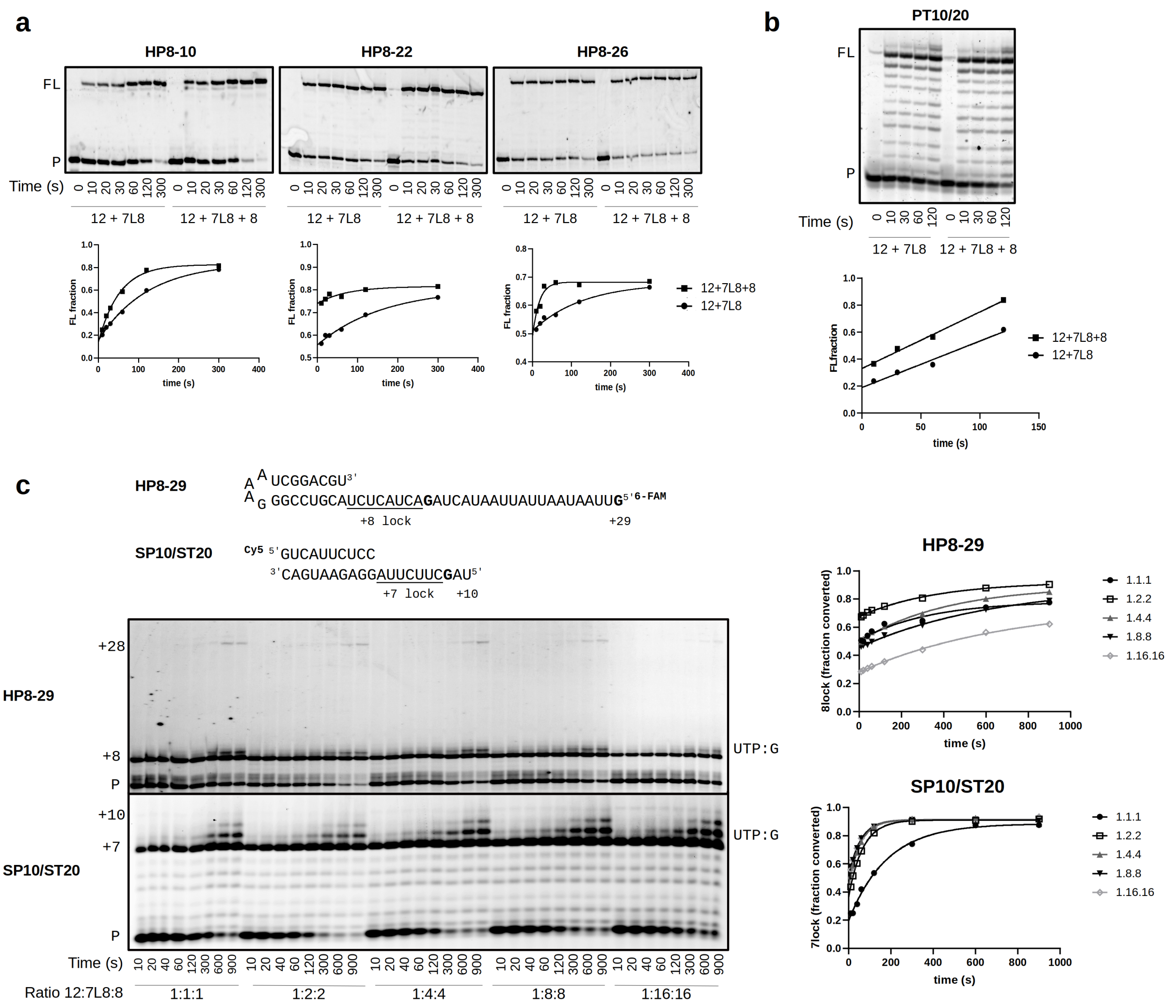


**Extended data Fig. 2. Comparison of activity of nsp12:7L8 and nsp12:7L8:8 complexes. a,** Extension of varied HP and **b,** PT substrate by either the nsp12+nsp7L8 complex (1:10 molar ratio) or nsp12+7L8+8 complex (1:10:5 molar ratio). Graphs show the fraction of full-length (FL) product produced at each timepoint relative to the total RNA. **c,** Extension of HP and PT substrates by the nsp12:7L8:8 complex at different molar ratios (as indicated). Reactions were run with run with 50 µM final concentration of ATP, UTP and GTP (omitting CTP) to form +8 and +7 stalled complexes for the HP and PT substrates respectively. Read-through products resulting from a nucleotide UTP misincorporation event against the templated G are seen for both substrates (UTP:G). Graphs show the fraction of stalled +7/+8 product relative to the amount of total RNA.

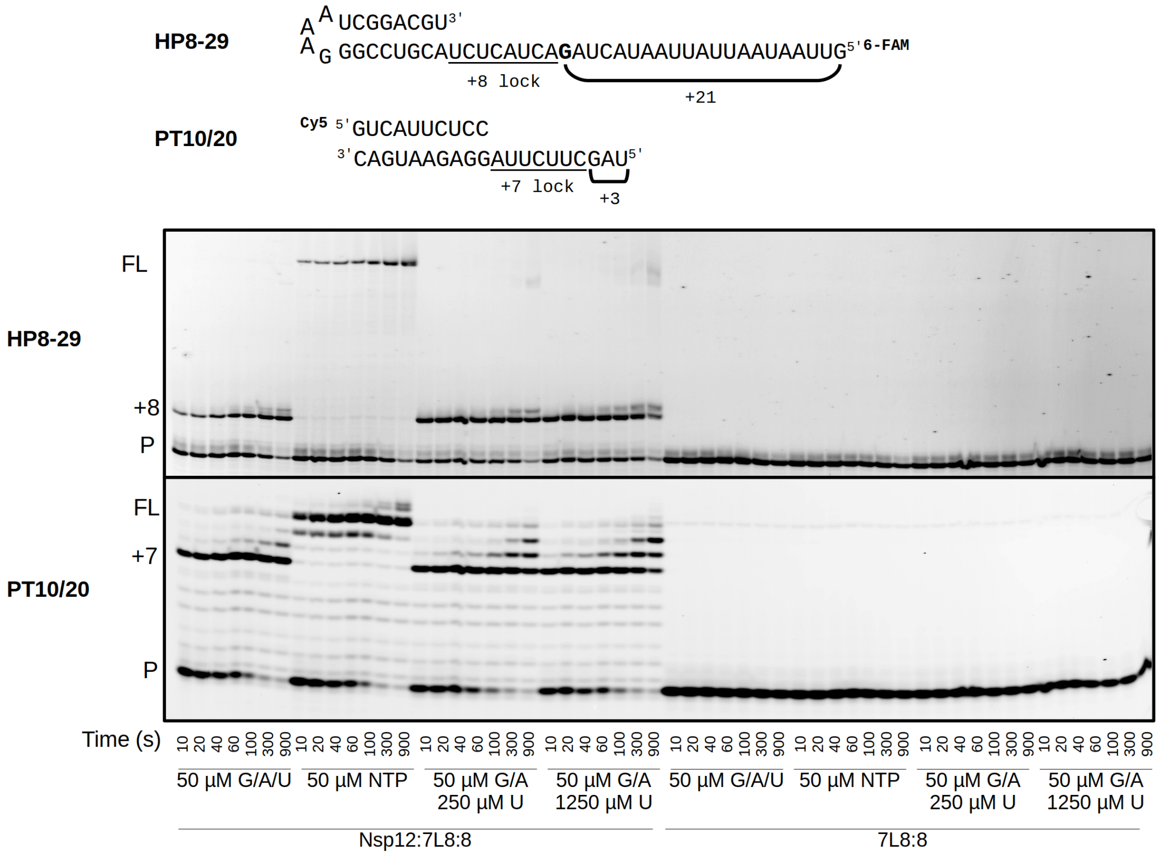


**Extended data Fig. 3. Elongation with varied NTP combinations in the presence and absence of nsp12.** Extension of HP and PT substrates by either the nsp12:7L8:8 complex or with nsp7L8:8 alone. Reactions were initiated with varied NTP conditions, either all four NTPs, or with GTP, ATP and UTP only (omitting CTP), with UTP concentration varied 5-fold (50, 250 and 1250 µM as indicated). Nsp8 has previously been reported to contain primase-like noncanonical RdRp activities[41-43] but under our conditions no primer elongation is observed in the absence of nsp12.

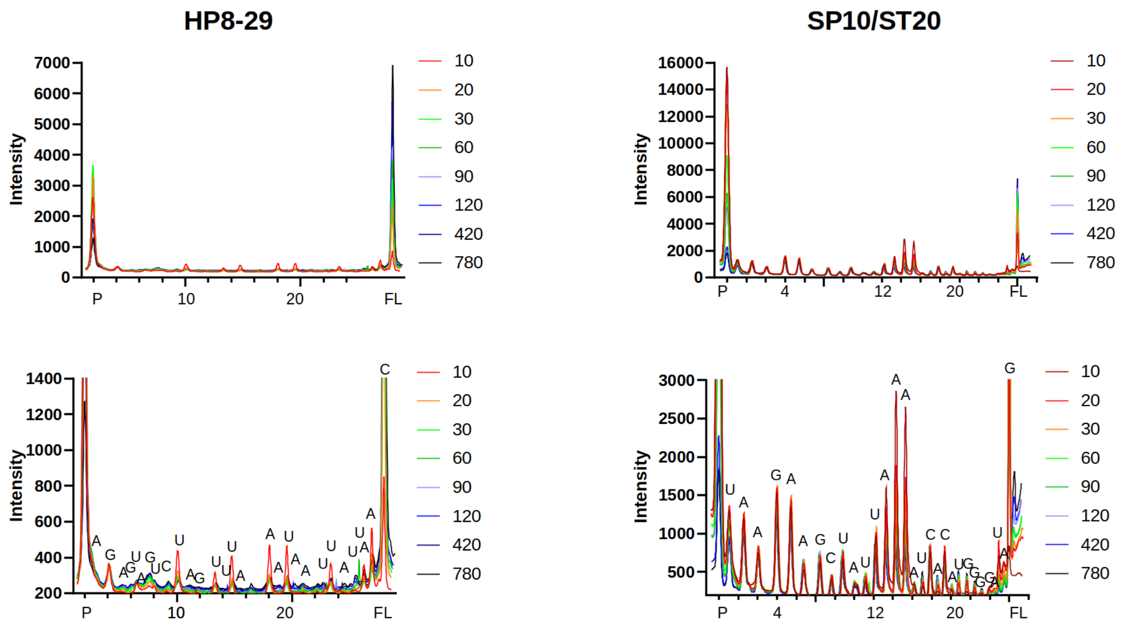


**Extended data Fig. 4. Intermediate species over time shown in Fig. 1.** Lane profiles of manual quench experiments performed with HP8-29 and PT10/40 substrates were obtained using ImageJ (Fiji). Peaks were normalized along the x-axis, and are shown from primer (P) to full-length (FL) for each substrate. Top panels show raw lane intensity for each substrate. Bottom panels are scaled 5X for relative peak analysis, with band assignments shown. Colors represent data taken from each timepoint in seconds.

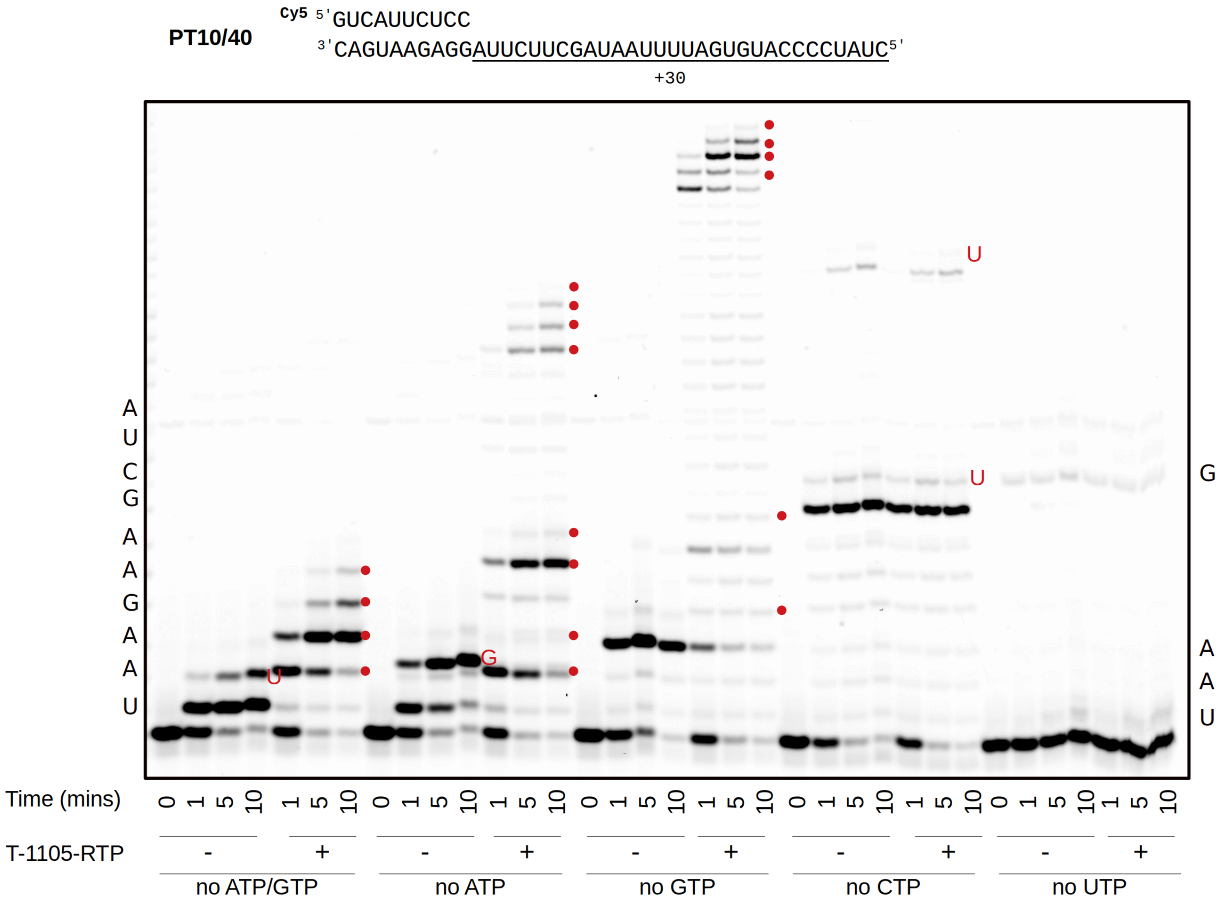


**Extended data Fig. 5. Nsp12:7L8:8 incorporation of nucleoside analogues into RNA.** Extension of PT10/40 substrate in the presence or absence of T-1105-RTP (10 µM) with either one or two NTPs omitted and 50 µM of other NTPs. Incorporation of analogues shown with red dots. Nucleotides on the side of gel indicate correct nucleotide incorporation for first 10 positions, and nucleotides in red on gel indicate incorporation of incorrect nucleotides.

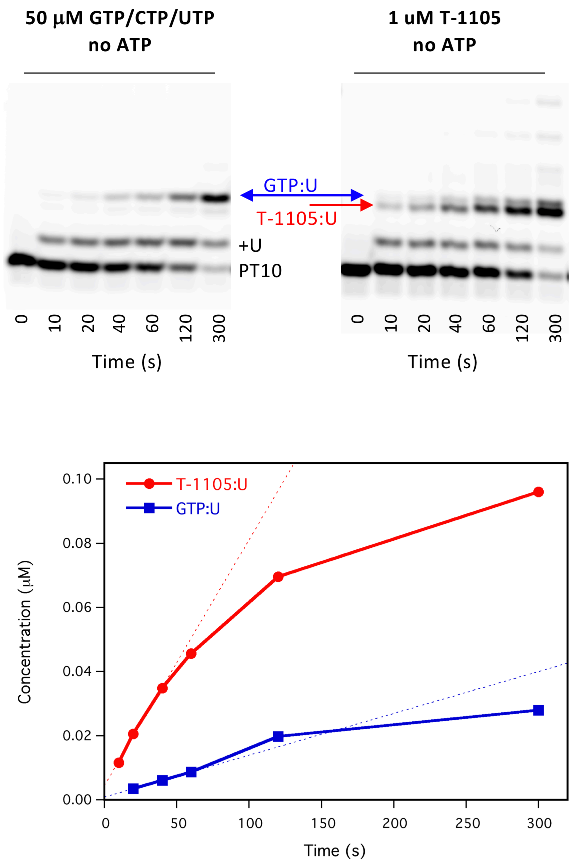


| | T-1105:U | GTP:U | Ratio |
|---|---|---|---|
| Intercept (nM) | 4.5±1.2 | 0.88±0.04 | 5±1 |
| Slope (nM/s) | 0.77±0.05 | 0.13±0.08 | 5±3 |

**Extended data Fig. 6. Comparison of natural GTP:U mismatch and T-1105-RTP:U incorporation levels.** Elongation reactions with the PT substrate done in the absence of ATP with 50 µM each GTP, UTP, and CTP show rapid addition of the first uracil followed by slow misincorporation of a GTP:U mismatch. In the presence of 1 µM T-1105 (with 50 µM each GTP, UTP, and CTP) the analogue incorporation is on a similar timescale as the native GTP:U mismatch. Analysis of incorporation rates show a burst phase followed by linear product buildup over a 60-second timeframe that is consistent with the measured lifetime of the nsp12-7L8-8 elongation complex. Both the burst amplitude and linear rate indicate that 1 µM T-1105 is incorporated ~5-fold faster than the natural GTP:U mismatch. Correcting for the concentration difference, the T-1105 reaction is estimated to be ~250-fold more efficient than a natural GTP:U mismatch.

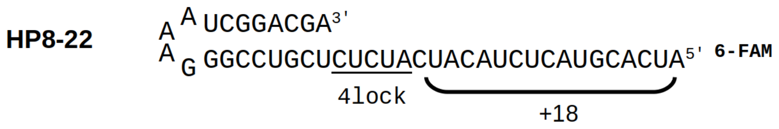


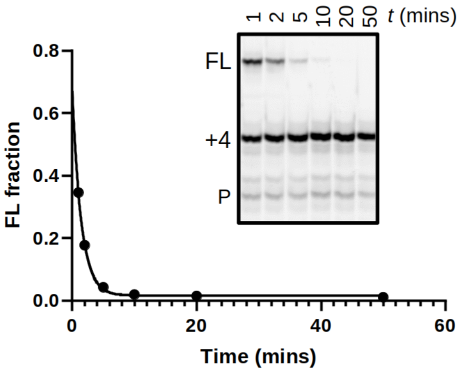


| 12:7L8:8 | Half Life (mins) | 95% CI |
|---|---|---|
| **1.1.1** | 1.003 | 0.8492 to 1.189 |
| **1.2.2** | 1.092 | 1.000 to 1.192 |
| **1.4.4** | 0.9025 | 0.7738 to 1.057 |
| **1.8.8** | 1.066 | 0.9872 to 1.153 |
| **1.10.10** | 0.9558 | 0.8336 to 1.099 |
| **1.10.5** | 0.9816 | 0.8568 to 1.128 |

**Extended data Fig. 7. Formation of a stable elongation complex on HP RNA.** Nsp12:7L8:8 complexes at varied ratios were incubated with HP RNA and 50 µM GTP and ATP for 5 mins to promote formation of 4nt-lock, and diluted 1:2 in high salt to prevent RNA rebinding. Reactions were chased at indicated timepoints with 50 µM all NTPs. Chase reactions were quenched after 30 sec and analyzed by gel electrophoresis (1:4:4 ratio shown). Stability half-life was calculated from the ratio of full-length (FL) product produced relative to the total amount of +4-lock + FL product at each timepoint. Graph shows stability data for 1:4:4 complex only, with calculated stability for all ratios shown in the table.

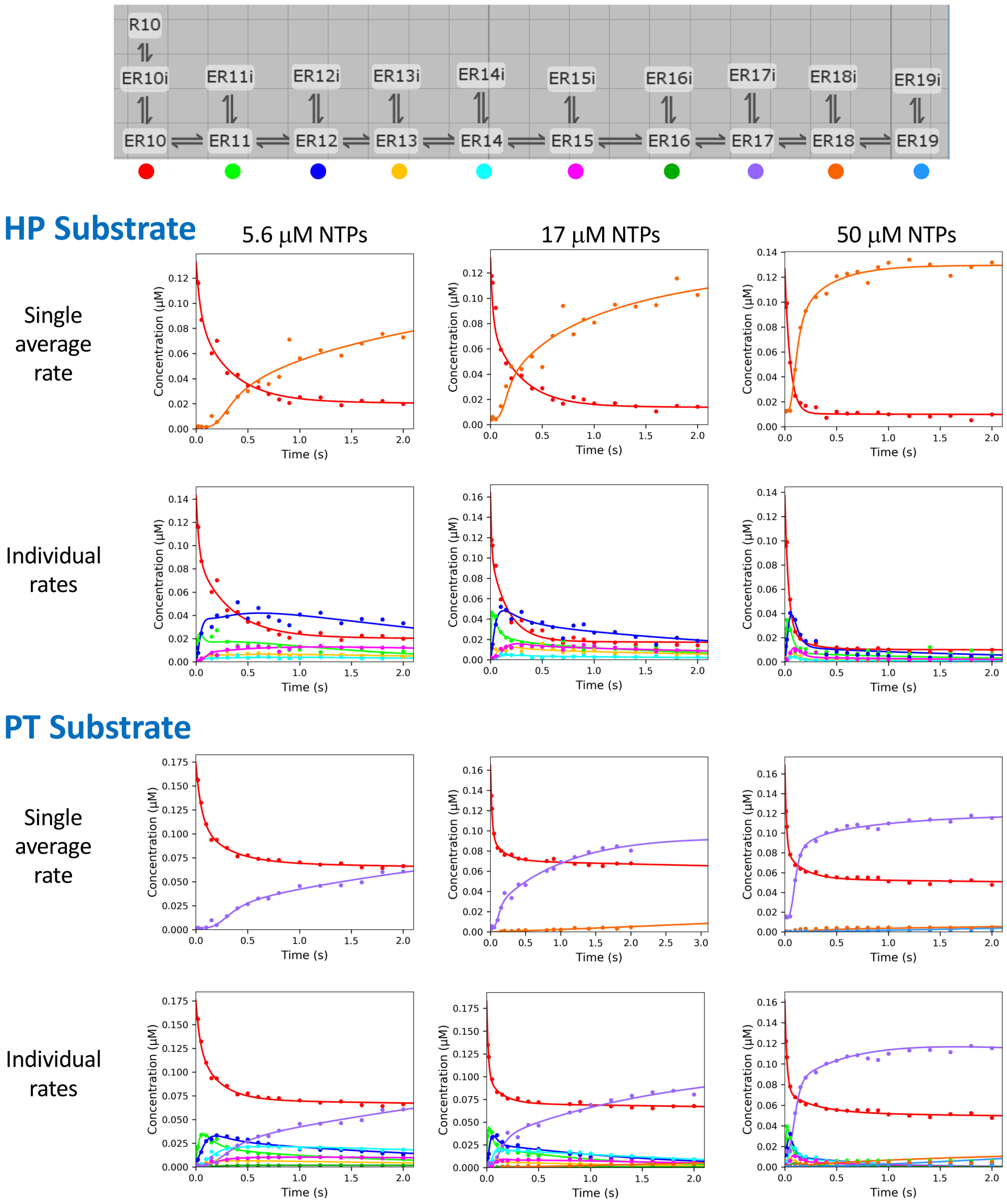


**Extended data Fig. 8.** Kinetic modeling of pre-steady state quench-flow data. Elongation kinetics were analyzed by data fitting using KinTek Explorer with a model for sequential addition steps wherein in every elongation state was in an equilibrium with an off-pathway inactive species to account for the large amounts of intermediate species observed in the data. Initial concentrations of the ER10 species reflect pre-assembled polymerase-RNA complexes capable of immediate burst phase elongation, and the R10 species reflects unbound RNA that is slowly depleted in the reaction, leading to the slow final phase of full-length product formation. An average rate was determined by making all elongation steps equal and considering only the primer and full-length product concentration data, while the individual rates analysis allowed each nucleotide addition step to vary independently. The inactivation/reactivation steps were always fit to a single common set of rates for all species. See Table S1 for complete results from the kinetic fitting.

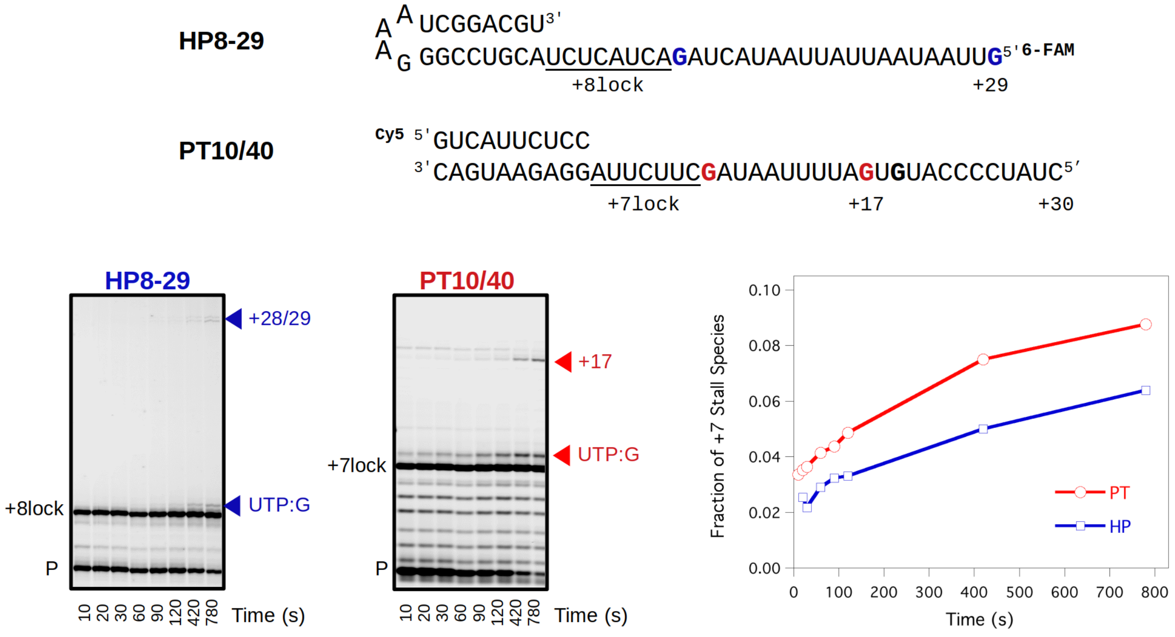


**Extended data Fig. 9. Efficiency of bypassing the no-CTP stall site on PT and HP templates.** For each timepoint, the total intensity of all bands greater than the +7 (PT) or +8 (HP) stall species was divided by the intensity of the stalled species to calculate an overall bypass efficiency on each substrate. Averaged across all timepoints, the ratio of these efficiencies shows PT complex is 1.45±0.11-fold more prone to bypass than the HP complex

| NTP: Template Pairing | **PT** 5uM NTP | **PT** 16 uM NTP | **PT** 50 uM NTP | NTP: Template Pairing | **HP** 5 uM NTP | **HP** 16 uM NTP | **HP** 50 uM NTP |
|---|---|---|---|---|---|---|---|
| **Single Average Rate Analysis** | | | | | | | |
| Avg $k_{pol}$ | 18–20<br>18.7±0.9 | 45–51<br>48±2 | 64–68<br>66.0±1.3 | Avg $k_{pol}$ | 21–28<br>23±3 | 40–55<br>46±6 | 90–103<br>95±8 |
| ER→ERi | 3.8–4.8<br>4.2±0.4 | 8.8–11<br>9.7±0.8 | 3.1–3.9<br>3.5±0.2 | ER→ERi | 2.6–5.0<br>3.5±0.9 | 3.9–6.9<br>4.9±1.5 | 4.2–7.4<br>5.3±1.5 |
| ER←ERi | 0.5–0.7<br>0.60±0.09 | 1.7–2.1<br>1.83±0.14 | 0.75–1.2<br>0.93±0.17 | ER←ERi | 0.5–1.0<br>0.8±0.2 | 0.9–1.7<br>1.2±0.4 | 3.1–5.1<br>3.9±1.0 |
| **UTP** | – | 0.16–0.25<br>0.20±0.04 | 0.12–0.16<br>0.14±0.01 | | | | |
| **UTP:G vs Avg** | – | **1/240** | **1/470** | | | | |
| **Individual Incorporation Rates Analysis** | | | | | | | |
| UTP:A | 25.0±1.0 | 65±2 | 51.0±1.2 | ATP:U | 38±3 | 104±8 | 29.4±1.3 |
| ATP:U | 12.6±0.7 | 34.0±1.6 | 39.8±1.4 | GTP:C | 37±6 | 23.8±1.7 | 46±4 |
| ATP:U | 8.7±0.7 | 22.2±1.1 | 39.9±1.5 | ATP:U | 9.0±1.5 | 8.9±0.6 | 25±2 |
| GTP:C | 57.6±0.8 | 84±9 | 151±15 | GTP:C | 53±12 | 30±3 | 140±40 |
| ATP:U | 12.4±1.7 | 20.6±1.1 | 45±2 | UTP:A | 80±30 | 80±20 | 230±110 |
| ATP:U | 21±3 | 39±3 | 70±5 | ATP:U | 21±4 | 19±2 | 76±13 |
| GTP:C | 120±40 | 280±130 | 210±40 | GTP:C | Not fit | Not fit | Not fit |
| **UTP:G** | – | **0.21±0.04** | **0.23±0.03** | UTP:A | Not fit | Not fit | Not fit |
| UTP:A | – | | 1.6±0.8 | | | | |
| **UTP:G vs UTP:A** | – | **1/310** | **1/220** | | | | |
| ER→ERi | 6.5±1.1 | 8.0±0.6 | 3.9±0.3 | ER→ERi | 8±2 | 3.2±0.5 | 2.6±0.5 |
| ER←ERi | 0.63±0.06 | 1.2±0.1 | 1.7±0.2 | ER←ERi | 0.8±0.2 | 0.57±0.13 | 0.4±0.2 |

**Extended data Table 1. Pre steady-state elongation rates from kinetics modeling of EDTA quench-flow data.** Values obtained from fitting with KinTek Explorer using the kinetic model shown in figure S8. The single average rate values are listed both 90% confidence intervals and optimal values with standard errors. The +7 GTP:C and +8 UTP:A steps were not included for the HP template because the observed bands were very weak and could not be definitively identified when peak fitting the gel lane profiles. Values in red represent putative UTP:G mismatch incorporation events observed on the PT substrate and the ratios of these rates to either the single average rate or to a cognate UTP:A addition.
